# Large-scale expansions of Friedreich’s ataxia GAA•TTC repeats in human cells are prevented by LNA-DNA oligonucleotides and PNA oligomers

**DOI:** 10.1101/2022.07.04.498742

**Authors:** Anastasia Rastokina, Negin Mozafari, Jorge Cebrián, C.I Edvard Smith, Sergei M. Mirkin, Rula Zain

**Author notes:** To whom correspondence may be addressed: Rula Zain. Shared first author (Anastasia Rastokina, Negin Mozafari). Shared last author (Sergei M. Mirkin, Rula Zain).

## Abstract

The human disease Friedreich’s ataxia (FRDA) is caused by expansions of GAA•TTC repeats in the first intron intron of the frataxin (*FXN)* gene, and both intergenerational and somatic expansions are crucial for disease development. We and others have shown earlier that expanded GAA•TTC repeats can form an intramolecular triplex structure (H-DNA). Here we studied the effects of locked nucleic acid (LNA)-DNA mixmer oligonucleotides and peptide nucleic acid (PNA) oligomers on the expansion of GAA•TTC repeats in cultured human cells. Our experimental system employes a mammalian/yeast shuttle plasmid containing a selectable cassette to detect repeat expansions. Using our in-house *in vitro* triplex-specific DNA cleavage assay, we first confirmed H-DNA formation by the (GAA)_100_•(TTC)_100_ repeat in the selectable cassette and demonstrated that the designed LNA-DNA oligonucleotides as well as PNA oligomers are able to disrupt this structure. We then found that both LNA-DNA mixmers and PNA oligomers prevent repeat expansions in human cells. In the accompanying paper, we show that expansions of GAA•TTC repeats in this experimental system occur during replication fork stalling, regression and restart at the repetitive run. We hypothesize, therefore, that triplex DNA formation by the GAA•TTC repeats is a key to their instability, while LNA-DNA oligonucleotides and PNA oligomers counteract repeat expansions by disrupting the triplex at the fork or preventing triplex formation upon fork reversal.

## Introduction

Friedreich’s ataxia (FRDA) is an autosomal recessive disorder with an incidence of 1 in 50,000 in European and American populations ^1,2^. It was first recognized by the German physician Nikolaus Friedreich in 1877 ^3^. FRDA is caused by an expansion of the GAA•TTC trinucleotide repeat located in the first intron of the *FXN* gene ^4^. The majority of FRDA patients have two expanded GAA•TTC alleles, while a minority have an inactivating single nucleotide polymorphism (SNP) in one allele and expansion in the other, disrupting the formation of an intact frataxin protein, is a less frequent cause of FRDA ^5^. Pathogenic expanded alleles consist of around 66-1700 repeats, while the normal allele contains from 7 to 22 GAA repeats ^6,7^. The number of the repeats is inversely related to the age of onset and the severity of disease ^8^. Unfortunately, we are still lacking FRDA therapeutics and only symptomatic management strategies are available.

The nuclear *FXN* gene encodes the protein frataxin, which predominantly localizes in mitochondria but is also expressed weakly in nuclei, endoplasmic reticulum and microsomes ^9,10^. The expansion of GAA•TTC repeats in the *FXN* gene results in reduced expression of frataxin mRNA and protein. Frataxin is responsible for iron-sulfur (Fe-S) cluster biosynthesis and diminished levels of frataxin lead to mitochondrial iron accumulation and elevation of cellular oxidative stress ^11-13^. The role of Fe-S clusters is diverse, from electron transfer and iron regulation to DNA repair. Dysregulation of frataxin due to reduced frataxin levels leads to Fe-S deficiency and increase of toxic iron in mitochondria and subsequent neurodegeneration ^13^. Neurodegeneration in FRDA is characterized by damage in the spinal cord, dorsal root ganglia (DRG) and cerebellum ^14,15^. Patients often develop sensory and motor dysfunction at puberty and eventually lose their ability to walk, becoming wheelchair-bound. They also suffer from progressive cardiomyopathy, which is the leading cause of death for this disease ^16^.

*FXN* downregulation is triggered by the formation of an intramolecular DNA triplex, known also as H-DNA, by the expanded GAA•TTC repeat, which impedes transcription elongation ^7,17,18^ ultimately leading to local heterochromatin formation and gene silencing ^19^. Two types of triplex structures can be formed by pathogenic expanded GAA•TTC repeats: pyrimidine-purine-pyrimidine (Y*RY) or purine-purine-pyrimidine (R*RY) where the third strand is homopyrimidine or homopurine, respectively ^20^. Chemical probing of expanded GAA•TTC repeats in supercoiled plasmids has shown that Y*RY triplex is the prevalent conformation *in vitro* ^21-23^.

The GAA•TTC repeats are unstable during germline transmission from parent to offspring, which can result in large-scale repeat expansions or contractions between generations ^24,25^. Pathogenic GAA•TTC repeats also tend to expand further in somatic cells resulting in disease progression during the affected individual’s lifespan ^26-28^. The most prominent expansions occur in DRG followed by cerebellum and heart ^29^, which could likely explain the late onset of cardiomyopathy compared to the earlier pathology seen in DRG ^29^.

Mechanisms responsible for GAA•TTC repeat expansions were primarily studied in model experimental systems. First, these repeats were shown to stall replication fork progression in bacteria ^27^, yeast ^30^, mammalian cell culture ^31,32^ and patient iPSC cells ^33^.

Further, studies in yeast have demonstrated that expansions and contractions of GAA•TTC repeats occur at some point during DNA replication ^34-38^. Replication was also implicated in the stability of GAA•TTC repeats within the SV40-based mammalian episome ^39^.

Other machinery implicated in GAA•TTC repeat instability is DNA mismatch repair (MMR). In yeast, MMR, and specifically the MutLα complex appears to cleave H-DNA, which results in chromosomal fragility in dividing cells ^40^. Notably, most affected cells in FRDA are post-mitotic, thus, repeat instability in those cells is independent of DNA replication. In an experimental yeast system to study GAA•TTC repeat instability in non-dividing, quiescent cells, MMR counteracted repeat expansions by triggering the formation of deletions or gene conversion events ^41^. Contrasting results were obtained in a human cell line characterized by progressive, small-scale expansions of GAA•TTC repeats: They appear to be independent of cell division and promoted by the mismatch repair complex MutLγ ^42^. Similarly, MMR promoted GAA·TTC repeat expansions in iPSCs derived from FRDA patient fibroblasts ^43^. Expansions of GAA•TTC repeats were also studied in humanized mice. In this system, intergenerational expansions of GAA repeats were inhibited by a mismatch repair system, while somatic expansions in the cerebellum and DRG were promoted by MMR ^44,45^.

The accompanying paper describes a new experimental system to simultaneously follow replication fork progression through GAA•TTC repeats and their propensity to expand in human cell culture. The progression of the replication fork through the repeat was analyzed via electrophoretic analysis of plasmid replication intermediates isolated from human cells, while repeat expansions that occurred in human cells were detected upon plasmid transformation into yeast. We found that repeat expansion events likely occur during replication fork regression within the repeat and its subsequent restart.

In this study, using a triplex-specific cleavage assay, we show that an H-DNA structure is efficiently formed by the (GAA)_100_•(TTC) _100_ repeat in the plasmids used in our replication/expansion studies in human cells. We have previously reported that chemically modified sequence-specific oligonucleotides disrupt H-DNA formation by GAA•TTC repeats in supercoiled plasmids. Here, we examined the ability of those oligonucleotides to affect the instability and expansion of GAA•TTC repeats in human cells. We found that the LNA-DNA mixmers, hereafter referred to as LNA-ONs, and the corresponding PNA oligomers dramatically reduce the expansion frequency of GAA•TTC repeats in human cells. These data hold promise for the development of these compounds for the treatment of FRDA, which is currently incurable.

## Results

### LNA ONs bind and disrupt the intramolecular triplex formed by the (GAA)_100_•(TTC)_100_ repeat in the experimental plasmid for detection of repeat expansions in human cells

The accompanying paper describes our experimental system to study GAA repeat expansions in human cells. It is based on the mammalian-yeast shuttle vector, pJC_GAA100 (Fig, 1A). In brief, this 12 kb-long plasmid contains a fragment of the SV40 virus carrying Tag and viral replication origin for its replication in human cell, a yeast ARS-CEN element, which makes it a single-copy yeast plasmid, and a UR-(GAA)_100_-A3 selectable cassette for the detection of repeat expansions.

**Figure 1.**
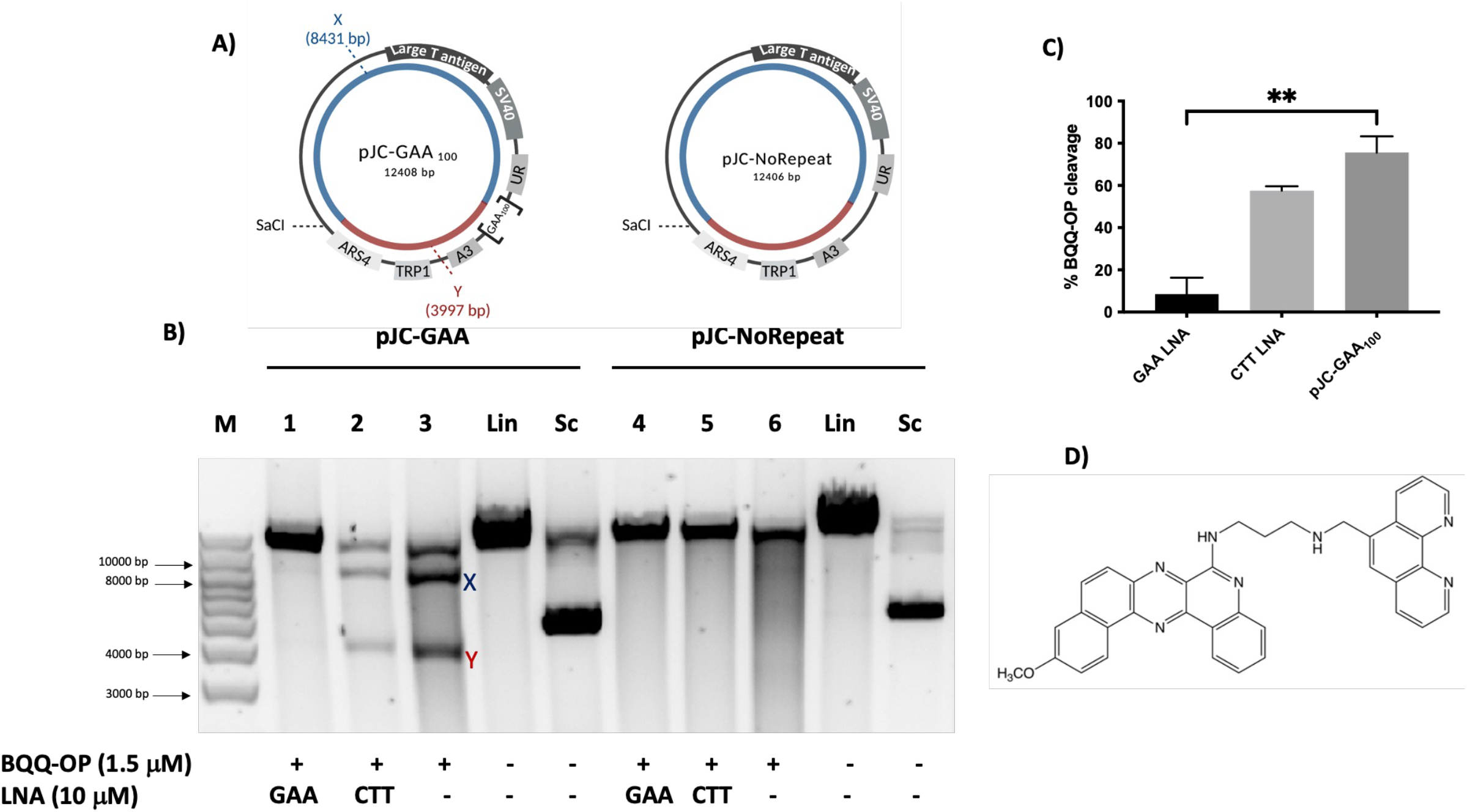
BQQ-OP mediated DNA cleavage of H-DNA-forming (GAA)_100_ repeats in the presence of LNA oligomers. **A)** Schematic illustration of pJC-GAA and pJC-NoRepeat. The H-DNA forming site in pJC-GAA is indicated as (GAA)_100_ and there is no H-DNA forming site in pJC-NoRepeat sequence. The two DNA fragments generated by BQQ-OP (benzoquinoquinoxaline 1,10-*ortho*-phenanthroline) triplex cleavage followed by unique site restriction digestion are indicated as X (8431 bp) and Y (3997 bp) in pJC-GAA and the same reaction would result in a linearized fragment only in pJC-NoRepeat. **B)** Representative agarose gel for pJC-GAA and pJC-NoRepeat incubated with 10 μM GAA(LNA-DNA) mixmer (lanes 1 and 4 respectively), or CTT (LNA-DNA) mixmer (lanes 2 and 5 respectively) or in the absence of LNAs (lanes 3 and 6 respectively). BQQ-OP-mediated triplex specific cleavage of pJC-GAA and pJC-NoRepeat was performed in the presence of Cu^2+^ and 3-mercaptopropionic acid (MPA) followed by unique site restriction digestion with SacI. As controls, supercoiled (Sc) and linearized (Lin) variants of both plasmids and molecular weight DNA ladder (M) are shown. **C)** Graph showing the percentage of BQQ-OP-mediated triplex specific cleavage of pJC-GAA in the presence of GAA and CTT (LNA-DNA) mixmers or in the absence of LNA-DNA mixmers (pJC-GAA). The values represent the ratio between the intensity of DNA double strand cleavage (X+Y) to the total band intensity of the particular lane and are shown as mean with S.D. (n=2). No cleavage was obtained in pJC-NoRepeat and not included in the graph. ** indicate P≤ 0.01 compared to the plasmid in the absence of LNA oligomers. **D)** Chemical structure of benzoquinoquinoxaline 1,10-*ortho*-phenanthroline (BQQ-OP).

We have previously shown that LNA-modified ONs, which are complementary to either the CTT or GAA strand (CTT LNA-ONs and GAA LNA-ONs, respectively) interact differently with expanded (GAA)_n_•(TTC)_n_ repeats cloned from FRDA patient cells. The GAA LNA-ON significantly disrupted and abolished the pyrimidine-motif H-DNA (H-y DNA) ^22^ by forming a duplex-invasion complex, whereas the CTT LNA-ON targeted the GAA single strand in this particular triplex motif. These studies were performed with a plasmid that was much different from the pJC_GAA100 plasmid: it was shorter, contained longer repeats and additional flanking sequences from human DNA.

To assess the effect of LNA-ONs binding to the (GAA)_100_•(TTC)_100_ repeat in the pJC_GAA100 plasmid, we used the BQQ-OP triplex-specific cleavage assay. BQQ-OP (Figure 1) is a low-molecular weight compound that consists of a benzoquinoquinoxaline derivative (BQQ) conjugated to a 1,10-*ortho*-phenanthroline (OP). BQQ recognizes and intercalates into inter- and intramolecular triplex structures. We have previously reported the ability of BQQ-OP to bind to triplex structures formed by FRDA repeats and introduce double-strand DNA breaks ^22,23,46^.

Here, we performed the BQQ-OP cleavage assay to validate that the (GAA)_100_•(TTC)_100_ repeat in pJC_GAA100 forms a triplex and to evaluate the effect of LNA-ONs and PNA oligomers (Table 1) on the structure of the repeat.

**Table 1.**
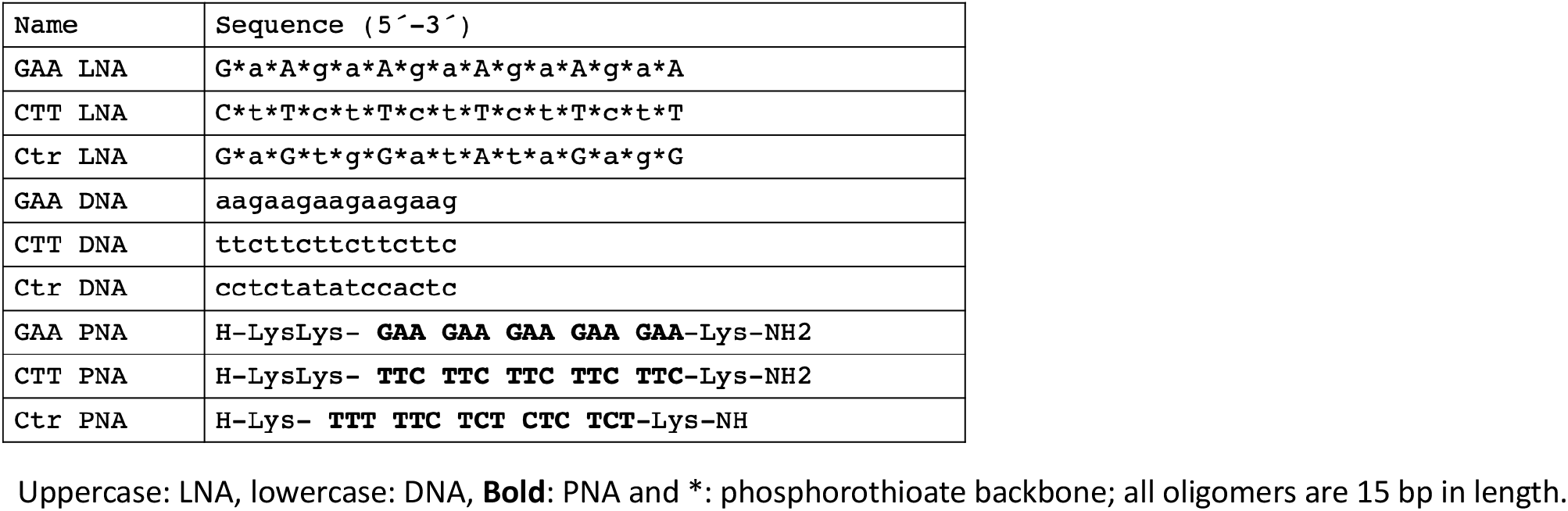
LNA-DNA mixmers and PNA oligonucleotides used in this study.

If the (GAA)_100_•(TTC)_100_ run in the supercoiled pJC_GAA plasmid forms H-DNA, BQQ-OP cleavage followed by restriction digestion with the *Sac*I restriction enzyme should produce two DNA fragments ∼8431 and 3997 bp-long, which are marked as X and Y in Figure 1A. Similar treatment of the control plasmid, which lacks the repeat sequence (pJC-Norepeat), should produce a single, 12,406 bp-long linear DNA fragment corresponding to the full-length plasmid cleaved by *Sac*I (Figure 1A). Indeed, the two expected DNA fragments were detected after BQQ-OP treatment of pJC_GAA, confirming H-DNA formation by the repeat (Figure 1B lane 3). At the same time, the control plasmid did not show any cleavage by the BQQ-OP (Figure 1B lane 6).

Incubation of the repeat-containing plasmid with GAA LNA-ON dramatically (∼10-fold) decreased BQQ-OP cleavage (Figure 1B lane 1). These data show that the GAA LNA-ONs practically obliterated H-DNA in accord with our previous findings ^22^. On the other hand, CTT LNA-ON only marginally affected the triplex-specific DNA cleavage (Figure 1B lane 2). Likewise, GAA PNA oligomers eradicate H-DNA in the repeat-containing plasmid while the CTT PNA oligomer has no such effect. (Supplementary Figure 1.)

### LNA-ONs inhibit GAA repeat expansions occurring during replication in human HEK-293T cells

To analyze expansion of the (GAA)_100_•(TTC)_100_ repeat in human cells, the pJC_GAA100 plasmid was transfected into HEK-293T cells, allowed to replicate for 48 hours, isolated, treated with the DpnI restriction enzyme to eliminate non-replicated bacterial DNA, and transformed into *S. cerevisiae*. The detection of repeat expansion events was possible due to the presence of the *UR*-(GAA)_100_-*A3 TRP1* selectable cassette containing the (GAA)_100_•(TTC)_100_ repeat in an intron of the artificially split *URA3* gene. Repeat expansions interrupt splicing of the *URA3* gene, making yeast resistant to 5-fluoroorotic acid (5-FOA), which allowed us to detect these events among Trp^+^ 5-FOA-resistant yeast transformants. The expansion baseline was defined by transforming the pJC_GAA100 plasmid isolated directly from bacteria, which did not undergo replication in mammalian cells, into yeast. In all cases, PCR analysis was performed to confirm repeat expansions in 5-FOA-r transformants. The frequency of repeat expansions was estimated as the ratio of PCR-confirmed 5-FOA-r clones to the total number of Trp^+^ clones using FluCalc software ^47^.

Data in Figure 2 demonstrate that the frequency of the (GAA)_100_•(TTC)_100_ repeat expansions increased ∼5-fold over the baseline upon plasmid replication in HEK-293T cells (P = 0.0043). To examine the effect of chemically modified oligonucleotides that disrupt H-DNA on repeat expansions in human cells, HEK-293T cells transfected with the pJC_GAA100 plasmid were treated with two different LNA-ONs: GAA LNA-ON or CTT LNA-ON described above. Strikingly, both oligonucleotides reduced the frequency of repeat expansions practically down to the baseline level, while a control oligonucleotide, which does not bind to the target sequence, had no effect (Figure 2). We conclude, therefore, that repeat-specific LNAs prevent GAA•TTC repeat expansions in human cells.

**Figure 2.**
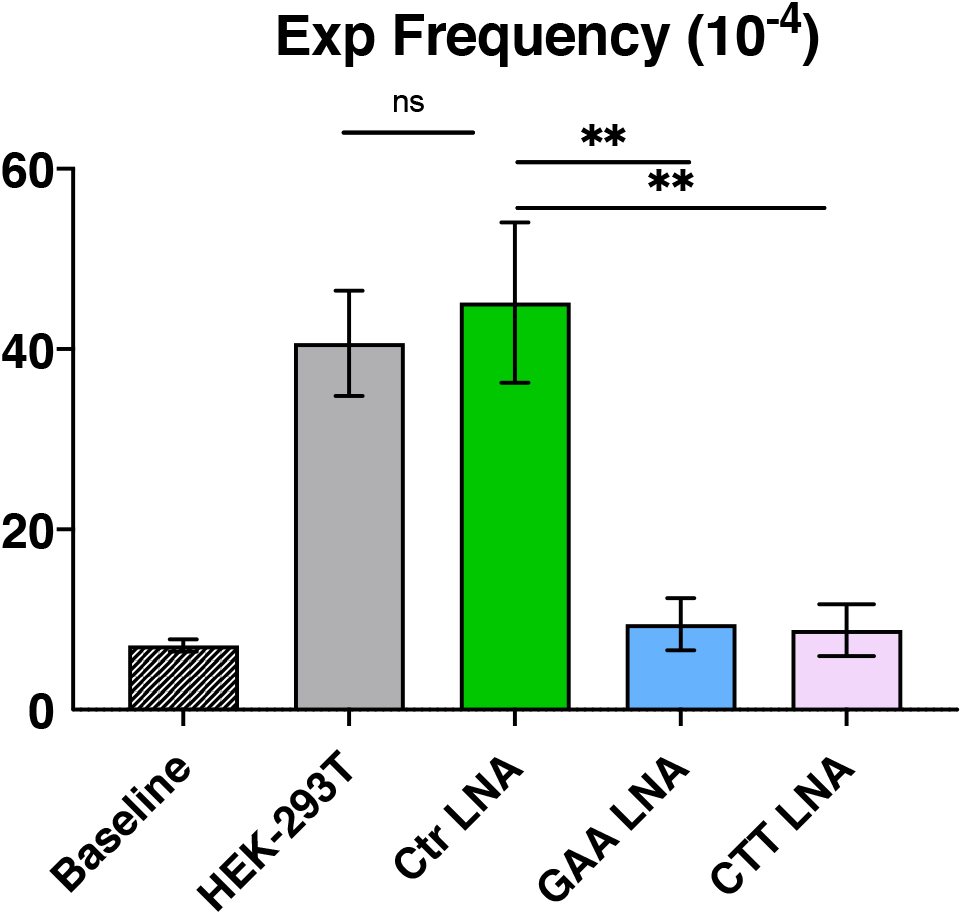
LNA-DNA mixmers inhibit GAA•TTC repeat expansions occurring during replication in human HEK-293T cells. Baseline represents the frequency of the (GAA)_100_ repeat expansion after *E. coli* plasmid transformation into yeast. Other bars in the chart show the frequency of GAA•TTC repeat expansions occurring during plasmid replication in HEK-293T cells in the presence or absence of various LNA-DNA mixmers. Error bars indicate standard error of the mean. Significance relative to the Scramble LNA-DNA mixmer control sequence frequency value was determined using a two-way Welch ANOVA test. ‘*’ *P* ≤ 0.05.

### Inhibition of GAA repeat expansions in HEK-293T cells by PNAs

To examine whether binding of another class of modified oligomers to the (GAA)_100_•(TTC)_100_ repeat would also affect its expansion frequency in human cells, we studied peptide nucleic acid (PNA) oligomers designed to bind this repeat. PNAs are DNA mimic oligomers with a pseudopeptide backbone ^48^ that are capable of binding and invading dsDNA in a sequence-specific manner and with high affinity owing to the lack of phosphate repulsion ^49^. We have previously studied the molecular interaction and binding mode of GAA- or CTT-containing PNAs (Table 1, GAA PNA and CTT PNA, respectively) to expanded GAA•TTC repeats in plasmid DNA *in vitro* ^22^. It was found that GAA PNA binds via a duplex-invasion mechanism and completely preclude H-DNA formation by the expanded repeats. CTT PNA, on the other hand, formed either a triplex-invasion complex or a Watson-Crick duplex when binding to the complementary polypurine strand of the DNA duplex ^22^.

Using the same experimental setting as in the previous section, HEK-293T cells carrying the replicating pJC_GAA100 episome were treated with either GAA PNA or CTT PNA. This was followed by plasmid DNA isolation and transformation into yeast to detect the frequency of expansion events that accumulated in human cells. Figure 3 shows that similarly to LNAs, both PNA oligomers reduced repeat expansions in human cells to the baseline level, while a control PNA oligomer, which does not bind to the target sequence had only a modest (1.5x) inhibitory effect. We conclude, therefore, that repeat-specific PNAs abolish expansion of the GAA•TTC repeat during replication in human cells.

**Figure 3.**
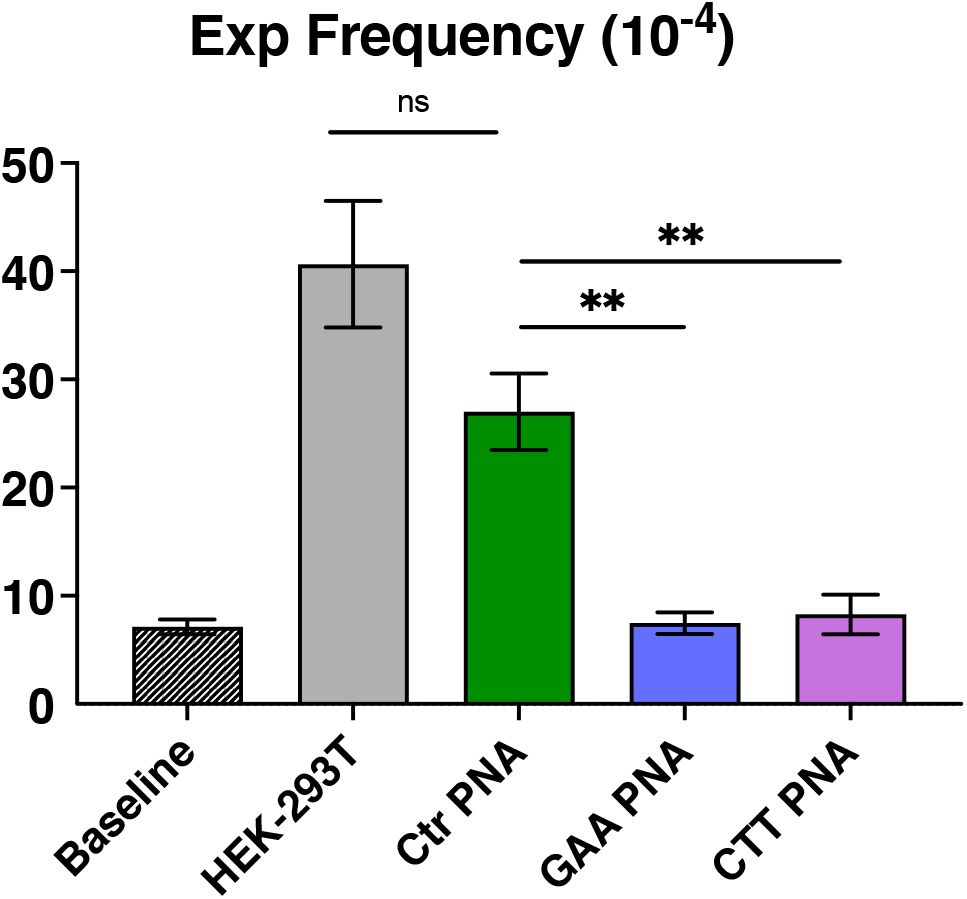
Inhibition of GAA•TTC repeat expansions occurring during replication in human HEK-293T cells. Baseline represents the frequency of the (GAA)_100_ repeat expansion after *E. coli* plasmid transformation into yeast. Other columns show the frequency of GAA repeat expansions that occurred during plasmid replication in HEK-293T cells in the presence or absence of various PNAs. Error bars indicate standard error of the mean. Significance relative to the Scramble PNA sequence frequency value was determined using a two-way Welch ANOVA test. ‘*’ *P* ≤ 0.005.

### Dose dependent inhibition of GAA repeat expansions in human HEK-293T cells by BQQ

In an accompanying paper, we have found that unwinding a triplex structure formed by the GAA•TTC repeat by the DDX11 helicase in the course of fork restart is needed for its expansion. We reasoned, therefore, that efficient stabilization of triplex DNA by chemical compounds might prevent expansions as well. We employed the benzoquinoquinoxaline derivative (BQQ) shown in Figure 4B to validate this hypothesis. We have previously reported on the efficiency of this heterocyclic compound to bind DNA polypurine•polypyrimidine sequences once they form an inter- or intramolecular triplex (H-DNA). BQQ intercalates in the triplex structures with its aminopropyl side chain positioned in the minor groove ^50,51^. BQQ has also been used to demonstrate the effect of H-DNA formation on transcription in a reporter model in bacteria ^52^ as well as triplex formation on a genomic level in mammalian cells ^53^ and its dissociation by ChlR1 helicase.

**Figure 4.**
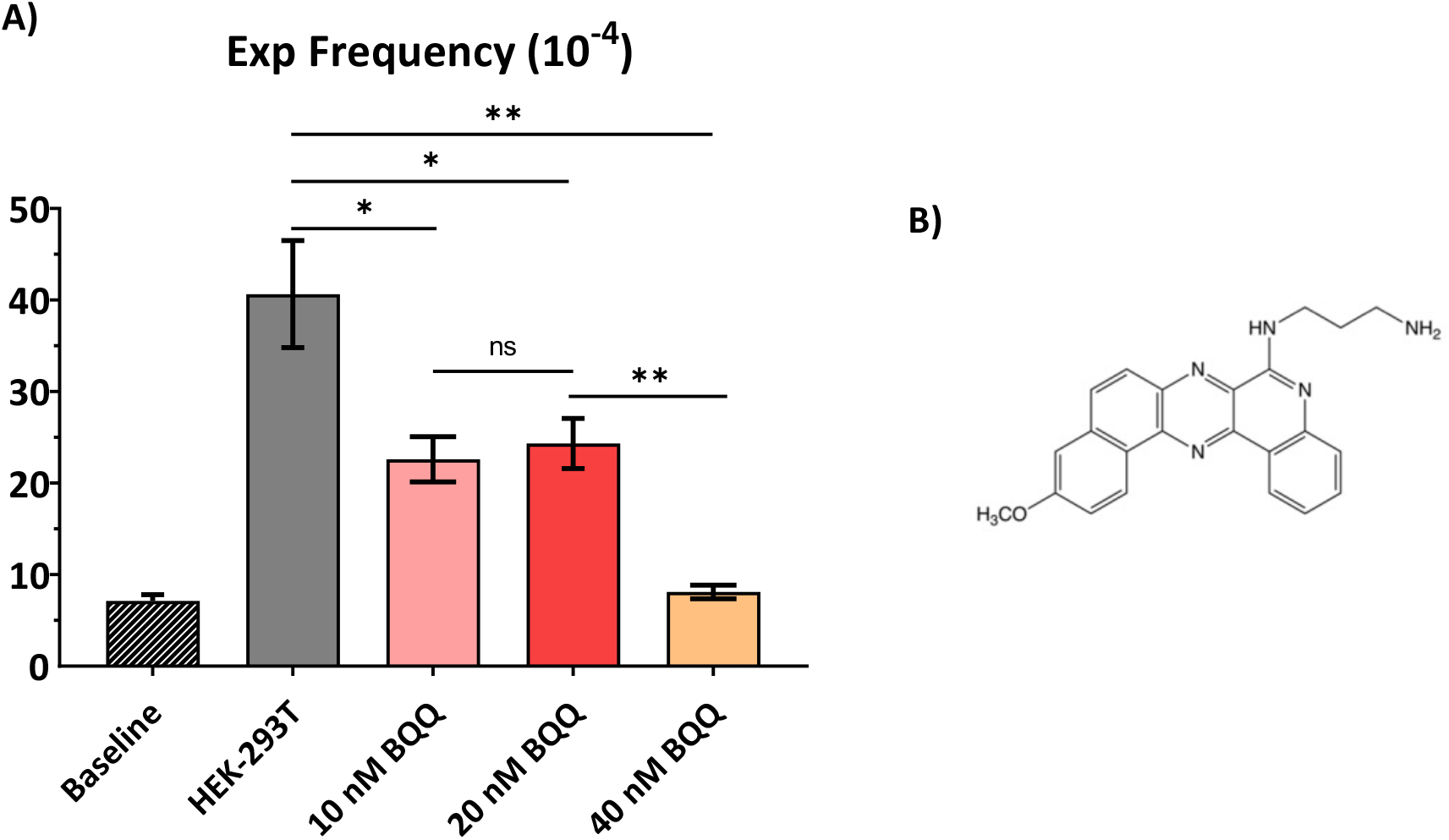
Dose-dependent inhibition of GAA•TTC repeat expansions in human HEK-293T cells by BQQ. Baseline represents the frequency of the (GAA)_100_ repeat expansion after plasmid transformation into yeast. Other columns show the frequency of (GAA)_100_ repeat expansions that occurred during plasmid replication in HEK-293T cells at various BQQ (benzoquinoquinoxaline) concentrations (10, 20 and 40 nM). Error bars indicate standard error of the mean. Significance relative to the HEK-293T frequency value was determined using a two-way Welch ANOVA test. (For 10 and 20 nM BQQ-OP concentrations *P* value is ≤ 0.05 for 40 nM *P*.≤ 0.005). **B)** Chemical structure of Benzoquinoquinoxaline derivative (BQQ).

We treated HEK-293T cells transfected with the pJC_GAA100 episome with BQQ at different concentrations. Figure 4A shows that BQQ treatment significantly decreased the frequency of the GAA•TTC repeat expansion as compared to untreated cells. The BQQ inhibitory effect was dose-dependent: ∼2-fold at 10 nM and 20 nM, and down to the baseline level at 40 nM. We conclude, therefore, that strong BQQ-mediated stabilization of the triplex formed by the (GAA)_100_•(TTC)_100_ repeat impedes repeat expansion in human cells.

## Discussion

Tandem-repeat instability is the cause of more than 50 neurodegenerative and neuromuscular diseases ^54^. Trinucleotide repeats (TNRs) form the largest group of disease-associated expanded repeats, and they are manifested in various hereditary diseases such as Fragile X syndrome ^55^, Huntington’s disease ^56^, numerous spinocerebellar ataxias ^57^, myotonic dystrophy type 1 ^58^ and Friedreich’s ataxia ^59^. Longer repeats are associated with a higher probability of expansions, which is also reflected in the severity and age of onset of the disease ^55^. TNRs can expand during both intergenerational transmission and within an individual’s somatic tissues. While the length of the inherited repeat is causative for disease development, its onset also depends on somatic expansions ^54^.

Several reports have demonstrated that expanded repeats can form alternative or non-B DNA structures, including hairpins ^60-64^, slipped-strand DNA ^65,66^, intramolecular triplexes (H-DNA) ^6,67,68^, quadruplex G4-DNA ^69-71^ or composite RNA-DNA structures, including R-loops ^72-74^.

The FRDA GAA•TTC repeat was shown to form an intramolecular triplex H-DNA conformation. This structure is formed upon strand dissociation in half the repeat and one of them folding back to form Hoogsteen or reverse-Hoogsteen hydrogen bonds with the remaining duplex. Either the polypurine (R) or polypyrimidine (Y) strand can be triplex-forming, while their complementary pyrimidine or purine strands, respectively, remain single-stranded ^21,75,76^. While H-y (Y*RY) triplex structures are usually pH dependent, for the GAA•TTC run, they are stable at neutral pH ^21,23^. Consequently, formation of both Py/Pu/Py and Pu/Pu/Py triplex was reported for this repeat at various ambient conditions in supercoiled DNA ^21,77-79^.

Using chemical and structural probing, our lab has previously established that the (GAA•TTC)_115_ repeat flanked by endogenous *FXN* gene sequences forms H-y triplex in supercoiled DNA. Furthermore, PNA oligomers or LNA-modified oligonucleotides were shown to invade the duplex and completely dissolve the preformed triplex structures ^22^.

Here we confirmed the formation of an intramolecular triplex by the (GAA•TTC)_100_ repeats in a plasmid designed to study repeat expansions in human cells using the BQQ-OP triplex-specific cleavage assay. We also report the distinctive behavior of GAA and CTT LNA-ONs on supercoiled plasmids containing 100 GAA repeats: GAA LNA-ONs abolished H-DNA formation, while CTT LNA-ONs had only a marginal effect. The difference between the current and previous CTT LNA-ONs data ^22^ could be due to the differences in DNA sequences flanking the GAA•TTC repeat, which might affect the nature and/or the stability of the triplex.

Numerous molecular mechanisms can contribute to the genetic instability of TNRs, involving DNA replication, DNA repair (MMR), and DNA recombination. The replication model for repeat instability stipulates the formation of alternative, dynamic DNA structures while the two DNA strands are being separated ^80^. It was shown that formation of these dynamic DNA structures during DNA replication impeded DNA polymerization, leading to DNA slippage and consequently repeat expansions or contractions ^81,82^. These dynamic DNA structures can also be additionally stabilized by the MutSβ MMR complex, which in turn, promotes repeat instability ^83^.

Here, we examined the effect of two distinct nucleic acids analogues, LNA and PNA, on the (GAA)_100._ expansion frequency. It appeared that both GAA and CTT oligomers containing either modification in their sequences dramatically decreased the expansion frequency in mammalian cells as compared to their corresponding controls.

This may be explained by their different mode of actions. GAA oligomers bind and disrupt H-DNA structure, which is assumed to play a role in DNA instability by interfering with DNA replication, transcription, and repair mechanisms ^84,85^. Two scenarios could be considered. First, GAA oligomers might disrupt H-DNA existing in a plasmid prior to DNA replication, a possibility discussed in ^31^ and (Figure 5). Alternatively, they can prevent formation of H-DNA during DNA replication either in front of the fork ^30^ or behind the fork during Okazaki fragment maturation ^86^. Our results in the accompanying paper implicate replication fork regression, caused by the unusual DNA structure of the GAA•TTC repeat, in its expansion in our experimental system. We hypothesize, therefore, that GAA LNA-ON and GAA PNA oligomers prevent the formation of H-DNA in front of the fork, thus, allowing replication to proceed smoothly (Figure 5).

**Figure 5.**
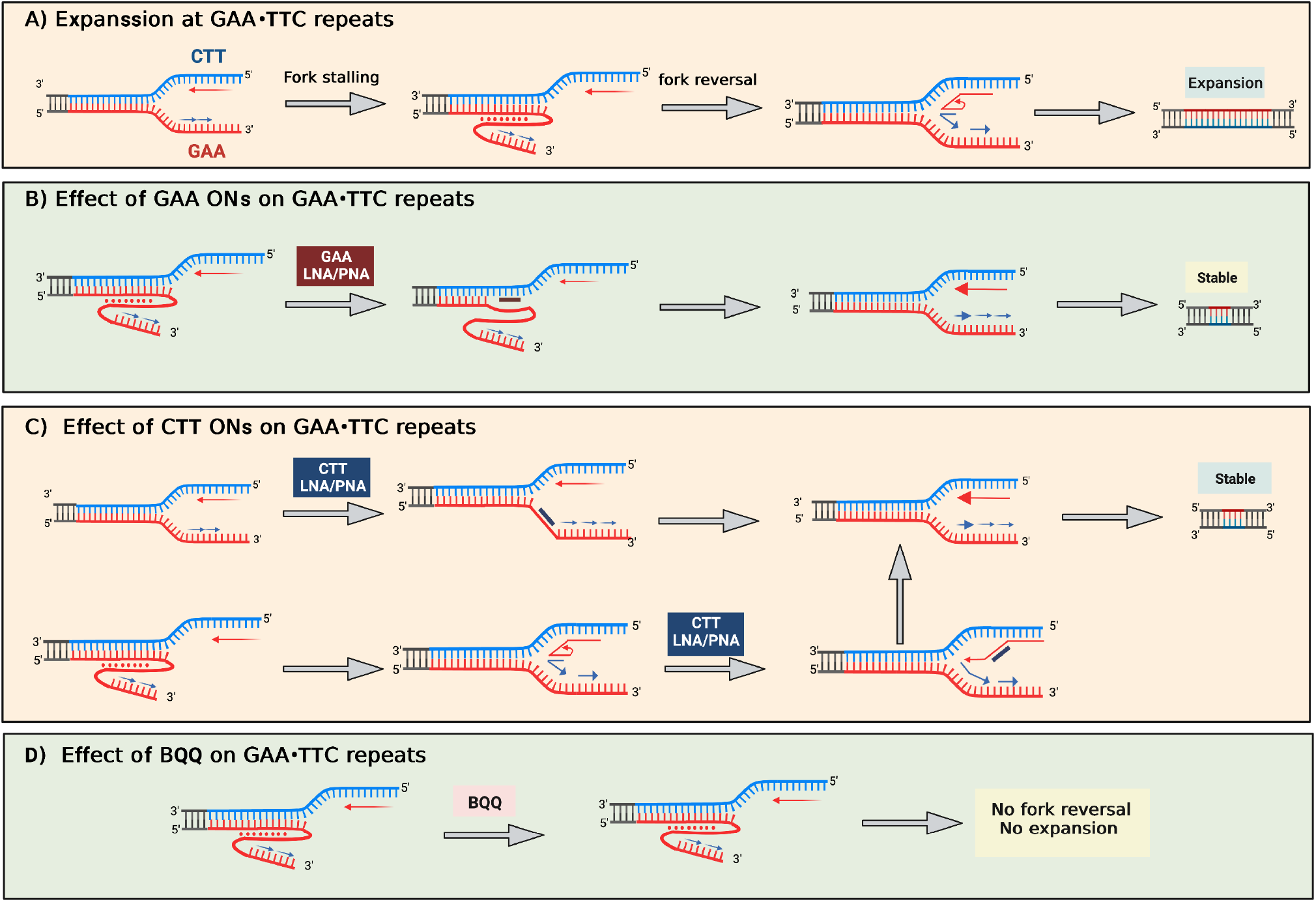
Proposed model for single strand ONs and oligomers (LNA-DNA mixmer or PNA) and BQQ preventing tandem-repeat expansion. GAA strand is shown in red and CTT strand is in blue. **A)** A model for (GAA•TTC) repeat expansion based on the formation of H-DNA during DNA replication and replication fork reversal. **B)** Prevention of GAA repeat expansion with GAA LNA/PNA by disruption of H-DNA before DNA replication (top) and by preventing the formation of H-DNA formation in front of the replication fork during replication (bottom). **C)** Prevention of (GAA•TTC) repeat expansion with CTT LNA/PNA blocking H-DNA formation in front of the replication fork during replication (top) and by blocking the formation of triplex in reversed fork (bottom). **D**) Prevention of H-DNA repeat expansion in the presence of BQQ stabilizing the H-DNA formed during replication and consequently prevention of replication fork reversal and, thus, expansion.

While CTT oligomers do not disrupt the preformed H-DNA structure, they also dramatically decrease expansion frequency. This makes the role of preformed H-DNA less likely. We could envision two scenarios. First, in order for the triplex to form ahead of the fork as proposed in ^87^, a portion of the lagging strand template containing (GAA)n run must fold back forming reverse Hoogsteen base pairs with the downstream duplex. CTT oligomers could prevent this from happening by occluding ssDNA with the GAA run (Figure 5). Secondly, our data in the accompanying paper suggest that formation of a triplex in the regressed fork prevents its proper restoration and leads to the triplex formation. An excess of CTT LNA-ONs and CTT PNA could prevent this from happening, thus, counteracting repeat expansions. It is important to note that both PNA oligomers and LNA-ONs have the distinct feature of being able to invade double-strand DNA. It is highly likely that this event is facilitated by the dynamics of DNA repeat regions forming non-B-DNA structures.

Finally, the specific triplex-stabilizing intercalating compound, BQQ, also decreases the frequency of GAA•TTC repeat expansions in a dose-dependent manner in our system. We hypothesize that this could be due to the excessive BQQ-mediated stabilization of a triplex ahead of the fork, making fork regression by DNA helicases more problematic (Figure 5).

Altogether, we report for the first time that chemically modified ss ONs and oligomers counteract the expansion of GAA•TTC repeats replicated in human HEK-293T cells. In FRDA patients, repeat expansions occur throughout life, significantly contributing to the disease severity and onset ^88^. Hence, the use of ONs could be a potential therapeutic strategy aimed at preventing somatic repeat expansions in FRDA patients.

## Methods

### Oligonucleotides

LNA and DNA oligonucleotides and PNA oligomers were purchased from Eurogentec S.A. The oligonucleotides and oligomers were purified with reversed phase HPLC, and quality control was performed with MALDI-TOF mass spectrometry. Control PNA was kindly provided by Prof. P. E: Nielsen, Department of Cellular and Molecular Medicine, University of Copenhagen ^48^.

### Plasmid

The plasmid pJC_GAA100 was constructed by conventional cloning methods in several steps using as backbone the pLM113 plasmid (gift from M. Lopes). First, pRS316 was digested by *Sal*I and *Sac*I to obtain the ARS4CEN6 sequence, which was then inserted between *Sal*I and *Sac*I of pML113 (gift from M. Lopes) creating a centromeric plasmid called pJW12 (8835 bp). The pJC_GAA100 plasmid (12408 bp) was obtained by inserting the *Ale*I-*Stu*I fragment of pYes3-T269-GAA100 plasmid, which contains UR-GAA100-A3 TRP1 cassette ^89^, into the blunt-ended *Eco*RV site of pJW12. In the resultant pJC_GAA100 plasmid, GAA repeats are on the lagging strand template for DNA replication from the SV40 origin. The pJC_GAA0 (No repeat) plasmid was obtained in a similar way except for inserting a *Ale*I-*Stu*I fragment from the pYES-TET644 [2]. All plasmid constructs were isolated from the *E. coli* SURE strain (Stratagene) and the integrity of (GAA)_100_ repeats was confirmed by Sanger sequencing. The pJC_TTC100 plasmid was obtained inserting the repeat-containing *Bsu36*I-*BstB*I fragment of pYes3-T269-TTC100 ^90^ into plasmid pJC_GAA100 using *Bs36*I and *BstB*I sites. All plasmid samples were isolated from the SURE strain (Stratagene) by the plasmid maxi-kit (Qiagen Inc.)

### BQQ-OP mediated DNA cleavage of H-DNA forming (GAA•TTC)_100_ repeats in the presence of LNA oligonucleotides

Plasmids pJC-GAA, pJC-TTC and pJC-NoRepeat (1 µg) containing 100 or no (GAA•TTC)_100_ repeats were incubated with 10 µM either GAA or CTT (LNA) oligomers (Table 1) in a buffer containing 10 mM sodium cacodylate, 100 mM NaCl and 2 mM MgCl_2_ at PH 7.5. As a control, plasmid with no oligonucleotide addition was treated in the same settings. The incubations were carried out at 37° C for 16 hours. BQQ-OP (1.5 µM) and CuSO_4_ (2.4 µM) were mixed and incubated at room temperature for 15 minutes. The BQQ-OP/ CuSO_4_ mixture was added to plasmid and oligonucleotide solutions and incubated at room temperature for 45 minutes. For the cleavage reaction to be initiated, 2 mM 3-mercaptopropionic acid (MPA) was added to the plasmid mixtures. The cleavage reaction was carried out for 3 hours at 37° C. The samples were purified with miniprep kit (Qiagen) and digested with 1U *Sac*I (Thermoscientific) for 1 hour at 37° C. The digested plasmids were analyzed with 0.7% gel electrophoresis with SYBR-gold (Invitrogen) staining. Gels were analyzed and quantified with VersaDoc MP system (Bio-Rad) and Quantity One software (Bio-Rad) respectively.

### Cell culture transfection

HEK-293T cells (ATCC) were grown in 10cm/10ml dish with the Dulbecco’s modified Eagle medium (DMEM) supplemented with 10% fetal bovine serum (FBS) and MycoZapTM Plus-CL (Lonza). Cells were transfected with pJC_GAA100 plasmid by using JetPRIME® (Polyplus-transfection) according to the manufacturer’s instructions. Briefly, cells were seeded on day 0 and co-transfected on day 1 withhttps://www.sciencedirect.com/topics/neuroscience/small-interfering-rna the GAA100 shuttle vector and ON of interest. After another 2 days, DNA was isolated to measure expansion frequencies.

### DNA isolation

Plasmid DNA was recovered 48 h post-transfection by a modified Qiagen Miniprep protocol as described ^91^. Briefly, cells were washed with PBS and then resuspended in Qiagen Buffer P1 and lysed in 0.66% sodium dodecyl sulphate 33 mM Tris-HCl, 6 mM EDTA, 66 μg/ml RNase followed by digestion with 0.5 mg/ml proteinase K for 90 min at 37 °C. Samples were subject to brief, 30 s, base extraction with 0.75 ml 0.1 M NaOH and proteins precipitated by addition of Qiagen Buffer P3 (4.2 M Guanidine-HCl, 0.9 M potassium acetate pH 4.8). Cell debris was pelleted at 29,000*g* for 45 min and supernatant loaded onto a Qiagen Miniprep spin column. Columns were washed with Qiagen Buffer PB (5 M Guanidine-HCl, 30% ethanol, adding 10 mM Tris-HCl pH 6.6) and 0.75 ml Qiagen Buffer PE (10 mM Tris-HCl pH 7.5, 80% ethanol) and plasmid DNA eluted using two volumes of 25 µl of Qiagen EB buffer.

### Yeast transformation

The plasmid DNA, prepared as described above, was digested for 1 h with 50 units of *Dpn*I (New England Biolabs) to eliminate unreplicated plasmids, EtOH precipitated, and resuspended in TE buffer. This DNA was used to transform yeast SMY537 strain (*MAT*a, *leu2-Δ1, trp1-Δ63, ura3–52, his3–200, bar1::HIS3*, can1::KanMX) ^89^ by lithium acetate method ^92^. A fraction of each transformation mixture (5% was plated onto SC-Trp plates (synthetic complete, lacking tryptophan), and the remaining 95% onto 5-FOA-Trp (synthetic complete, containing 0.09% 5-FOA and lacking tryptophan) to select for expansions. Colonies on each plate were counted after 3 days of growth at 30°C. The frequency of expansions (confirmed by PCR as described below) was determined as the number of colonies obtained on SC-Trp-FOA divided by the total number of transformants on SC-Trp, with appropriate correction for dilution factors. A critical concern with this assay is to ensure that the expansion events occurred in the tissue culture cells, and not during plasmid propagation in *E. coli*. We determined baseline expansion-events by transforming yeast cells directly with 100 ng of DNA plasmid preparation (without passing through mammalian cells). At least three independent transformants were tested to calculate the baseline and for each siRNA treatment.

### PCR analysis

To authenticate expansions and to determine their size, individual 5-FOA-resistant yeast colonies were disrupted with Lyticase as described in ^47^. Subsequent PCR amplification by Phusion Polymerase (ThermoFisher) used UC1 (5’-GGTCCCAATTCTGCAGATATCCATCACAC-3’) and UC6 (5’-GCAAGGAATGGTGCATGCTCGAT-3’) primers flanking the repeat tract for 35 cycles of 20 s at 98° C and 2 min 72° C with a final extension at 72° C for 4 min. The products were separated on 1.5% agarose gels. PCR product sizes were determined by comparison with 50 bp DNA ladder (New England Biolabs) using ImageLab™ software (Bio-Rad). PCR confirmation of expansions is especially important since randomly occurring mutations within the *URA3* gene itself could result in a 5-FOA-resistant phenotype.

### Statistical analysis

When indicated, statistical analysis was performed via Welch ANOVA test using GraphPad Prism version 8, GraphPad Software, San Diego, California USA.

## Funding

This study was supported by the grant R35GM130322 from NIGMS and by a generous contribution from the White family to S.M.M. Also, funding was kindly provided from Hjärnfonden, The Swedish Research Council, Swelife-Vinnova and CIMED and Region Stockholm (N.M, CIES, and RZ).

## Acknowledgments

We thank Catherine Freudenreich, Mitch McVey, and members of the Mirkin laboratory for their invaluable input on this project, Sanjukta Ghosh for her technical assistance and Julia Hisea for her terrific editorial help. We are also grateful to Prof. Peter E. Nielsen for providing the control PNA oligomer.

## Figure Legends

**Supplementary figure 1.**
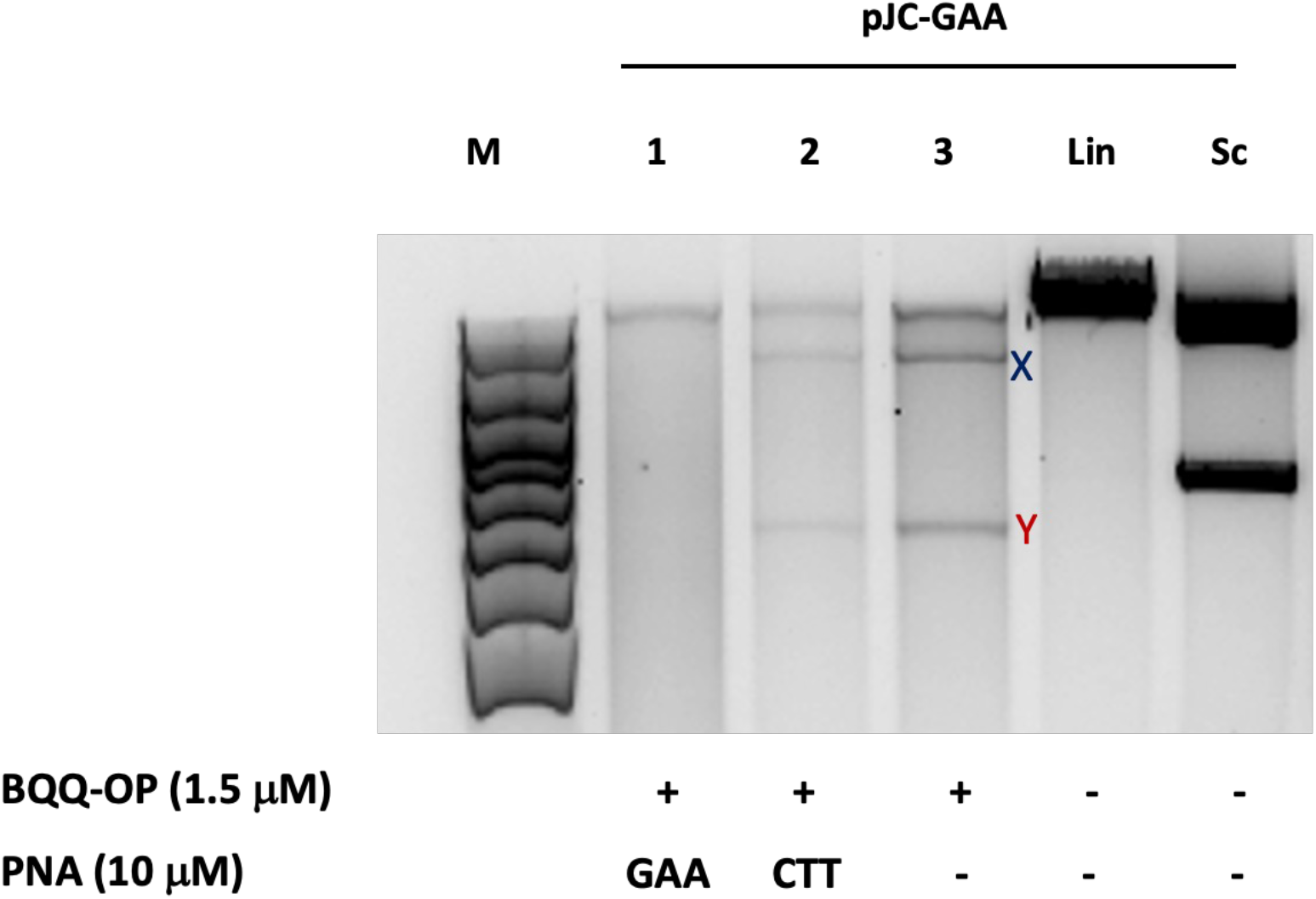
BQQ-OP mediated DNA cleavage of H-DNA forming (GAA)_100_ repeats in the presence of PNA oligomers. Agarose gel analysis for pJC-GAA plasmid incubated with 10 μM GAA PNA oligomer (lanes 1), or CTT PNA oligomer (lanes 2) or in the absence of PNAs (lanes 3). BQQ-OP-mediated triplex-specific cleavage of pJC-GAA was performed in the presence of Cu^2+^ and 3-mercaptopropionic acid (MPA) followed by unique site restriction digestion with SacI. As controls, supercoiled (Sc) and linearized (Lin) variants of plasmid and molecular weight DNA ladder (M) are shown.

## Notes

### Competing Interest Statement

The authors have declared no competing interest.

